# Single-Cell Analysis Reveals Critical Role of Macrophage Epsin in Regulating Origin of Foam Cell in Atherosclerosis

**DOI:** 10.1101/2024.08.29.610301

**Authors:** Kulandaisamy Arulsamy, Kui Cui, Xinlei Gao, Anna Voronova, Kaifu Chen, Hong Chen

## Abstract

Atherosclerosis is a chronic inflammatory condition characterized by the excessive accumulation of fat and lipid molecules, leading to the formation of foam cells and plaques in arterial walls. Dysfunction of vascular smooth muscle cells (VSMCs), fibroblast, endothelial cells, and macrophages is often associated with this pathology. We found that epsins accelerate atherosclerosis progression in individuals on a Western diet (WD). Using ApoE-deficient (ApoE^-/-^) and macrophage-specific epsin deletion in ApoE^-/-^ backgrounds (LysM-DKO/ApoE^-/-^) mice fed a WD for 16 weeks, we observed significantly reduced foam cell formation in LysM-DKO/ApoE^-/-^ mice compared to ApoE^-/-^ mice. Single-cell RNA sequencing identified 20 major cell types, including seven VSMC and five macrophage subtypes. Among the VSMC subtypes, modulating VSMC1 was involved in inflammation and migration, while modulating VSMC2 was associated with VSMC phenotype switching. In atherosclerotic mice, populations of modulating VSMC1, VSMC2, foamy-Trem2, and inflammatory macrophages increased, but significantly decreased in epsin-deficient mice. Modulating VSMC2 transition into macrophages occurred with a probability of 0.57 in ApoE^-/-^ mice, compared to 0.01 in LysM-DKO/ApoE^-/-^ mice. Epsin deletion also reversed endothelial dysfunction and downregulated cholesterol and glucose-mediated signals, as well as inflammatory ligands Il1b and C1qa. Our findings suggest that epsin deletion reduces foam cell formation and rewires VSMC and endothelial functions, offering a novel therapeutic strategy for atherosclerosis.

## Introduction

Atherosclerosis is a chronic inflammatory pathological condition in which the artery wall accumulates high deposits of fats, cholesterol, and other particles. This condition forms plaques that impede blood circulation and contribute to coronary heart disease, ischemic stroke, and abnormal aortic aneurysms^1–3^. Numerous serious risk factors, notably diet, are closely linked to the progression of atherosclerosis^4–6^. Specifically, a high-fat or Western diet positively correlates with pathological features such as arterial plaque formation^7,8^, extracellular matrix degradation^9,10^, loss of vascular smooth muscle cells^11,12^, endothelial cells dysfunction^13,14^ and immune cell infiltration and activation^15,16^. The inflammation observed in the arteries and atherosclerotic lesion formation arises from dysregulated communication between vascular smooth muscle cells, endothelial cells, and immune cells, particularly macrophages^17,18^. However, the molecular mechanisms underlying atherosclerosis progression in the context of a Western diet remain poorly understood.

Epsins comprise a family of endocytic adaptor proteins crucial for clathrin-mediated protein degradation^19–22^. Functionally redundant, epsin 1 and epsin 2 play pivotal roles in regulating lipid metabolism and exhibit specialized functions in modulating NOTCH, VEGFR3, and Wnt signaling pathways^23–26^. Over the past two decades, our group has extensively investigated the role of epsins in various contexts, including their involvement in embryonic development^23^, angiogenesis^24^, lymphomagenesis^25^, and lipid metabolism^27,28^. Additionally, we have identified epsins as critical regulators in tumor^25,29^ and atherosclerosis^30^ progression.

Furthermore, deletion of epsins in macrophages impedes the degradation of low-density lipoprotein receptor-related protein 1 (LRP-1), thereby enhancing efferocytosis and promoting an anti-inflammatory macrophage phenotype. This intervention reduces plaque formation by 50% in murine models^27^. Moreover, our recent findings demonstrate that macrophage-specific deletion of epsins in mice on a normal diet attenuates lipid uptake by downregulating CD36 levels and augmenting ABCG1-mediated cholesterol efflux^28^.

Despite notable advancements, a significant gap persists in our understanding of the molecular mechanisms driving atherosclerosis progression and its treatment on the Western diet. Epsin functions are predominantly associated with the progression of atherosclerosis ^27,30^. Therefore, formulating effective treatment strategies by targeting epsin is imperative. Here, we delineate key questions that remain unaddressed in current literature: i) What is the impact of the Western diet on the composition of cell types and their corresponding cell identity profiles in atherosclerosis, ii) How do cellular dynamics evolve during the progression of atherosclerosis, and what molecular events underlie these changes, iii) What are the intricate cell-cell communication signals that play pivotal roles in modulating cellular dynamics and inflammatory signaling pathways in the context of atherosclerosis, and iv) Which molecular targets are pivotal in modulating lipid metabolism and thereby influencing disease progression.

In this study, we aimed to address the questions mentioned earlier and elucidate the role of epsins in the progression of atherosclerosis. To achieve this, we utilized mouse models fed with the Western diet. Our findings revealed a notable reduction in the population of VSMCs and an increased prevalence of macrophages in atherosclerosis mice. We observed a predominant transition of VSMCs to a macrophage phenotype in WT/ApoE^-/-^, which was attenuated in epsin1 and 2 double knockout mice (LysM-DKO/ApoE^-/-^). In addition to this, the inflammatory signaling is downregulated after epsin deletion, suggesting their regulatory role in modulating inflammatory responses associated with atherosclerosis. Additionally, analysis of cell-cell communication pathways revealed a downregulation of cholesterol-mediated signaling, correlating with reduced foam cell formation and mitigated progression of atherosclerosis. Our study underscores the potential of targeted therapy directed at epsins as a promising avenue for the treatment of atherosclerosis.

## Material and Methods

### Animal models

All animal procedures were performed in compliance with institutional guidelines and mouse protocols approved by the Institutional Animal Care and Use Committee (IACUC) of Boston Children’s Hospital, MA, USA. Both male and female mice were used. C57BL/6 mice (stock #00664), ApoE^-/-^ mice (stock #002052) and LysM-Cre deleter mice (stock #004781) were purchased from Jackson Research Laboratory. Since epsins 1 and 2 (epsin1^-/-^; epsin2^-/-^) double knockout mice lead to embryonic lethality, we generated conditional Epsin1^fl/fl^; Epsin2^-/-^ mice previously described^28^. WT/ApoE^-/-^ and LysM-DKO/ApoE^-/-^ mice used in this study was generated in our previous study^28^. To generate atherosclerotic mouse model, WT/ApoE^-/-^ and LysM-DKO/ApoE^-/-^ mice were fed a Western diet (WD, Protein 17% kcal, Fat 40% kcal, Carbohydrate 43% kcal; D12079B, Research Diets, New Brunswick, USA) starting at the age of 6-8 weeks for 16 weeks.

### Single-cell RNA sequencing of the aorta cells through 10X genomics

WT/ApoE^-/-^ and LysM-DKO/ApoE^-/-^ mice were euthanized by CO2 inhalation. Then, we isolated aortas after 30 mL 1x PBS perfusion through left ventricular and quickly transferred to cold DMEM medium. Aortas from the two groups were minced into small pieces and digested with an enzyme solution (5mg/mL collagenase type I, 5mg/mL collagenase type IV, and 5mg/mL liberase) for 90min at 37 °C on a shaker. The cell suspension was filtered using a 40μm strainer and washed with PBS for two times. The cells were resuspended and ready for sequencing in PBS containing 0.04% bovine serum albumin, and their viability was over 90%.

Single-cell RNA-Seq library was performed using the protocol provided by 10X Genomics. In brief, the single-cell suspensions from both groups, reagents, gel beads and partitioning oil were loaded to 10X Chromium Chip G to generate single-cell Gel Beads-in-emulsion (GEMs, Single cell 3’ Reagent Kits v3.1, 10X Genomics). scRNA was barcoded through reverse transcription in individual GEMs followed by a post GEM-RT cleanup and cDNA amplification. Then, a 3’-gene expression library construction was performed. Finally, the library was sent for sequencing.

### Preprocessing of single-cell RNA sequencing reads

The raw sequencing reads in FASTQ format for every single cell was aligned to the mouse reference genome (mm10) using the cellranger software (version 6.1.2). It generated the gene expression count matrix, providing information on gene expression in raw counts for individual cells. Subsequently, the Seurat^31^ R package (version 4.1.0) was employed for downstream analysis. The “CreateSeuratObject” function was utilized to create the Seurat object from the gene expression count matrix, while the “PercentageFeatureSet” function computed the percentage of mitochondrial and ribosomal gene expression.

Subsequent filtering steps were implemented to remove low-quality cells based on the following criteria: i) cells expressing a number of genes between 200 and 5000, ii) cells with less than 10% mitochondrial gene expression, iii) cells with ribosomal gene expression below 20%, and iv) genes detected in fewer than 3 cells were considered rarely expressed and removed from the analysis. Additionally, mitochondrial genes and ribosomal protein-coding genes were excluded from the expression matrix before normalization. This stringent filtering process resulted in 8911 cells in the wild-type and 12342 cells in the epsin-deletion western-diet mice. To ensure a fair comparison between conditions, the number of cells in the epsin-deletion condition was down sampled to 8911 cells.

### Normalization, cell clustering and cell type annotation

For each condition, the gene expression count matrix underwent log-normalization using the “NormalizeData” function, followed by identifying highly variable genes with the “FindVariableFeatures” function. Subsequently, the “IntegrateData” function was employed to integrate the two conditions, and the integrated Seurat object was utilized for data scaling, principal component analysis (PCA), and UMAP visualization. Then, single-cell clustering was performed using the “FindClusters” function with a resolution parameter set to “0.8”. Marker genes were then identified using the “FindAllMarkers” function, which compares the expression profile of one cell cluster with the rest of all other cell clusters. During this step, parameters such as statistical tests were set to “wilcox”, the log fold change threshold value was set to “0.25”, and genes detected in at least “0.25” cells were considered. Furthermore, these cell clusters were annotated as cell types using the known marker genes in PanglaoDB^32^ and literature^33,34^.

### RNA velocity analysis

RNA velocity predicts the future states of cells by utilizing information on spliced and unspliced RNA content. To generate this, for each condition, we utilized the “bam” file containing read alignment information with the “velocyto run” function in the velocyto (version 0.17) command. Subsequently, the resulting spliced and unspliced information was used to identify the future states of cells using the default parameters in the scVelo^35^ Python package.

### Monocle trajectory analysis

For conducting the monocle^36^ trajectory analysis, we first converted the Seurat object into a monocle cell dataset object. Then, we used the “estimate_size_factors” function to calculate the size factors in the single-cell RNA-seq dataset. We retained the Seurat cell clusters and UMAP coordinates throughout the trajectory analysis. The root or starting cells of the trajectory were defined using the “order_cells” function. Moreover, the gene expression trends along the trajectory were visualized using the “plot_genes_in_pseudotime” function.

### SCIG uncovering the cell identity genes in a given cell type

Our newly developed SCIG method^37^ accurately identifies the complete spectrum of cell identity genes using RNA expression and genetic sequence profiles. We used the raw expression count of each gene as input to explore cell identity genes in the identified cell types. From the SCIG prediction result, we considered significant cell identity genes with P-values less than 0.05.

### NicheNet for inferencing the ligand-target mediated cell-cell communications

NicheNet^38^ (https://github.com/saeyslab/nichenetr) enables the exploration of ligand-target mediated cell-cell communications in the scRNA-seq dataset. We utilized the Seurat NicheNet wrapper function (nichenet_seuratobj_aggregate) with the following parameters: i) defining the sender and receiver cell populations, ii) conducting a differential expression test (log fold change cutoff set to 0.1 and genes expressed by at least 10% of cells) in the receiver cell population to identify enriched genes in ApoE^-/-^ and epsin-deletion western-diet mice, and iii) information of ligand-receptor and ligand-target gene regulatory networks. For the sender cell population, active ligands were determined based on their expression levels. The NicheNet analysis result was visualized by Circos plot which depicts the ligand-target gene interactions between the considered sender and receiver cell types.

### MEBOCOST for inferencing the metabolite-sensor mediated cell-cell communications

We utilized our Metabolic Cell-Cell Communication Modeling by Single Cell Transcriptome (MEBOCOST)^39^ algorithm (https://github.com/zhengrongbin/MEBOCOST) to predict cell-cell communications mediated by metabolites and sensor proteins at the single-cell level. Sender cells secrete or produce metabolites that travel and communicate with the sensor proteins of receiver cells. This algorithm first quantifies the presence of metabolites in every single cell using the expression levels of respective metabolite-producing and consuming enzymes. Subsequently, the metabolite abundance and expression levels of sensor proteins in each cell type are used to compute the communication score for each signal. During this analysis, we identified communication signals separately in both wild-type and epsin-deletion conditions and then combined them to identify differential communication signals. Flow plots were utilized to visualize the differentially regulated metabolite-sensor mediated signals in either the ApoE-deficient or epsin-deleted mice.

### Pathway enrichment analysis

We utilized the clusterProfiler^40^ R package to perform pathway enrichment analysis for marker genes, SCIG-derived cell identity genes, and differentially expressed genes. Functional pathways were obtained using the “enrichGO” function with the following parameters: organism database set to ‘mouse’, a significance threshold of P-value less than 0.05, and consideration of all gene ontology terms. Subsequently, these pathways were visualized using barplot, and cneplot function.

### RNA isolation, quantitative real-time PCR and RNA sequencing

Total RNA was extracted from aortas or primary macrophages with Qiagen RNeasy Mini Kit based on manufacturer’s instruction. The extracted RNA was used for qRT-PCR. For qRT-PCR, mRNA was reverse transcribed to cDNA with the iScript cDNA Synthesis Kit (Bio-Rad Laboratories, Inc., Hercules, CA, United States). 2 μL of the product was subjected to qRT-PCR in StepOnePlus Real-Time PCR System (Applied Biosystems) using SYBR Green PCR Master Mix reagent as the detector. PCR amplification was performed in triplicate on 96-well optical reaction plates and replicated in at least three independent experiments. The ΔΔCt method was used to analyze qPCR data. The Ct of β-actin cDNA was used to normalize all samples. Primers are listed in **Supplemental Table1**.

## Results

### Single-cell RNA sequencing reveals extensive VSMC heterogeneity and population changes between ApoE^-/-^ and epsin-deficient mice

To investigate the role of macrophage epsin in the Western diet, which is known to promote foam cell formation and atherosclerosis progression, we constructed Western-diet-fed mouse models of both ApoE-deficient and macrophage-specific deletion of epsin 1 and 2. Using these mouse models, we explored the cellular diversity of aortic cells through single-cell RNA sequencing **(Figure 1a)**. This approach yielded transcriptomic profiles of 29,724 and 11,414 cells in ApoE^-/-^ and LysM-DKO/ApoE^-/-^ conditions, respectively. We then rigorously filtered the data to ensure high-quality cells (n=8911 cells in each condition) for subsequent clustering and downstream analysis (**Figure S1a**). The Seurat single-cell clusters were annotated into 10 major cell types using known marker gene expressions, including vascular smooth muscle cells, endothelial cells, myofibroblasts, mesenchymal cells, macrophages, T cells, B cells, erythroid precursors, proliferating cells, and Schwann cells (**Figure S1b**). VSMC cells expressed typical markers such as Tagln, Acta2, and Myh11, while EC cells were identified based on marker genes like Pecam1, Cdh5, and Flt1. Macrophages were determined by the expression levels of Cd68, Lyz2, and Cd14 (**Figure S1c**).

**Figure 1:**
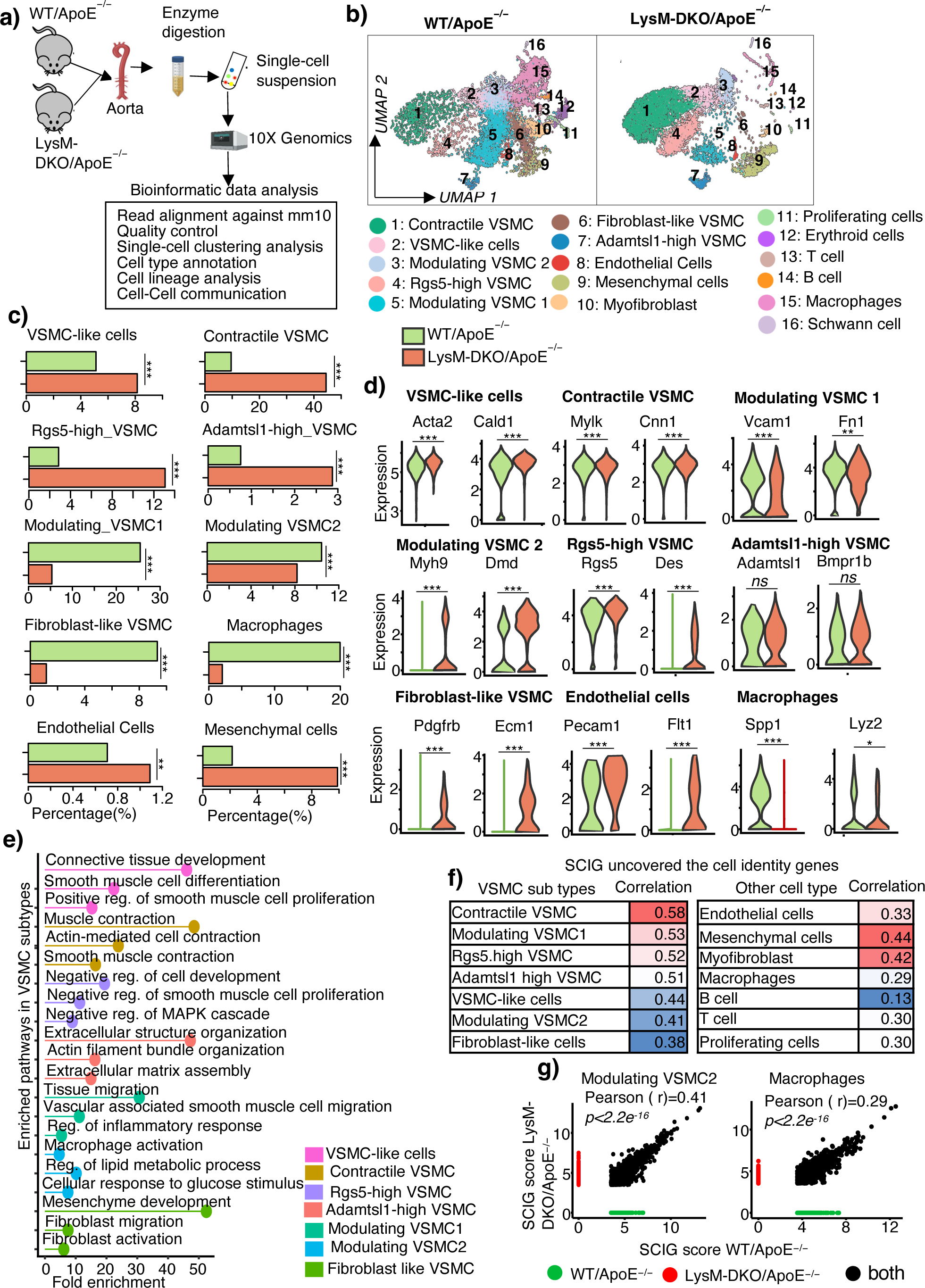
Single-cell RNA sequencing reveals novel cellular heterogeneity in western-diet-fed WT/ApoE^-/-^ and macrophage-specific epsin knockout mice. a) Workflow illustrating the preparation of aorta cells for single-cell RNA sequencing and subsequent bioinformatic data analysis. b) UMAP plot depicting 16 distinct cell types, including seven specific VSMC subtypes. c) Bar plot representing the proportions of each cell population in WT/ApoE^-/-^ and LysM-DKO/ApoE^-/-^ conditions. d) Violin plot showing marker genes used to annotate various VSMC subtypes, endothelial cells, and macrophages. e) Pathway enrichment analysis demonstrating the functional diversity of VSMC subtypes. f) The Pearson correlation between SCIG identified cell identity gene scores for each cell type between WT/ApoE^-/-^ and LysM-DKO/ApoE^-/-^. g) Scatter plot shows the observed correlation of cell identity genes in modulating VSMC2 and macrophages. *Note: P values determined by the Wilcoxon test. *, p-value < 0.05; **, p-value < 0.01; ***, p-value < 0.001*.

Further analysis of cell population sizes between ApoE^-/-^ and epsin-deficient mice revealed a decrease in populations of VSMCs, ECs, and mesenchymal cells in ApoE^-/-^ mice, contrasting with an increase observed in epsin-deficient mice (**Figure 1c**). This suggests a disruption in the physiological functions of VSMCs and ECs, particularly in the context of a Western diet. Remarkably, we observed a 10-fold decrease in the population size of macrophages in epsin-deficient mice (**Figure 1c**), indicating a potential protective role of epsins against foam cell formation and inflammatory responses. Additionally, immune cell populations, including B cells and T cells, were increased in ApoE^-/-^ mice (**Figure S1d**), suggesting their activation in the atherosclerosis progression. However, this phenomenon was reversed by the deletion of epsins in macrophages (**Figure S1d**). To validate these findings, we assessed the expression of the inflammatory marker gene TNF-⍺ using digested aortas from ApoE^-/-^ and LysM-DKO/ApoE^-/-^ mice. Consistent with our ScRNA data, we observed approximately a 10-fold increase in TNF-⍺ expression in ApoE^-/-^ mice compared to LysM-DKO/ApoE^-/-^ mice (**Figure S1e)**.

### Aorta of western diet mice hold diverse VSMC cellular and functional heterogeneity

It is known that VSMC cells have diverse subtypes, and each holds specific expression patterns and cellular functions^33,34^. Exploring the determining role of epsin in VSMC diversity, we specifically did a sub-clustering analysis for the VSMC population. We found seven different VSMC subtypes including Adamtsl1-high, contractile, fibroblast-like, modulating 1, modulating 2, Rgs5-high, and VSMC-like cells (**Figure 1b**). Adamtsl1-high VSMC cells express marker genes such as Adamtsl1, and Bmpr1b (**Figure 1d)**, which are involved in regulating extracellular matrix assembly (**Figure 1e**). Expression of genes Acta2, Cald1, Mylk and Cnn1 was observed in contractile VSMC and VSMC-like cells (**Figure 1d**), associated with smooth muscle cell contraction and functions (**Figure 1e**). Rgs5-high VSMC cells contain higher expression levels of Rgs5 along with Des genes (**Figure 1d**), playing a negative role in cell proliferation, consistent with previous findings^34^ (**Figure 1e**). The fibroblast-like VSMC subtype exhibits higher Pdgfrb, and Ecm1 gene expression (**Figure 1d**), implementing fibroblast cell-related functions (**Figure 1e**). Modulating VSMC1 shows enrichment in proliferation, migration, and adhesion by expressing Vcam1, and Fn1 (**Figure 1d and e**). Modulating VSMC2 cells express the Dmd, and Myh9 genes, enriched with a combination of VSMC migration, glucose, lipid metabolic process, and macrophage-related functions (**Figure 1d and e**). These results indicate that all VSMC subtypes express conventional marker genes such as Tagln, Acta2, and Myh11, suggesting a common VSMC identity (**Figure S1c**). However, they exhibit significant cellular and functional heterogeneity within their subtypes (**Figure 1d and e**).

Further, we compared the population sizes of each cell type between atherosclerosis and epsin-deficient mice. The contractile VSMC, VSMC-like cells, Adamtsl1-high VSMC, Rgs5-high VSMC were decreased and increased in ApoE^-/-^ and epsin-deficient mice, respectively (**Figure 1c**). Additionally, qRT-PCR was employed to validate the scRNA-seq findings. The results showed that Vcam1 and Myh9 were dramatically downregulated, while Rgs5 was significantly upregulated in LysM-DKO/ApoE-/- mice compared to the ApoE-/- mice (**Figure S2a-c)**. This indicates that the ApoE^-/-^ affects crucial VSMC functions such as muscle contraction, extracellular matrix assembly, and inhibition of VSMC proliferation. However, our target is that epsin deletion specifically in macrophages significantly alters these mechanisms. On the other hand, the modulating VSMC 1, modulating VSMC 2, and fibroblast-like VSMC increased in ApoE^-/-^ mice and its potential to increase inflammation and migration in VSMC cells. However, this impairment was rewired by epsin-deficient mice through control of inflammation and regulating physiological VSMC identity and functions (**Figure 1c**).

### Exploring the macrophage cell heterogeneity in the western diet of ApoE^-/-^ and LysM-DKO/ApoE^-/-^ mice

Elevation of foam cell formation and inflammatory pathways contribute to the progression of atherosclerosis^12,27^. To further characterize these phenomena in the macrophage epsin context, the sub-clustering analysis was conducted specifically on the macrophage population. Using marker gene expressions, we annotated five distinct macrophage subtypes: VSMC-derived, fibroblast-derived, inflammatory, foamy Trem2, and resident-like macrophages, as depicted in **Figure S3a**. Foamy Trem2 macrophages exhibited elevated expression levels of Trem2 and Abcg1, while inflammatory macrophages showed high expression of Tnf and II1b. Resident-like macrophages were identified by the expression level of Lyve1 and Mrc1 (**Figure S3b**). These were validated by profiling the expression level of marker genes through qRT-PCR (**Figure S3c**). Interestingly, a comparison of cell type population size between ApoE^-/-^ and epsin depleted mice revealed a significant decrease, ranging from 3-10-fold, in foam cell, inflammatory, and macrophages derived from VSMCs and fibroblasts cells (**Figure S3d**). However, the population of resident-like macrophage cells increased in LysM-DKO/ApoE^-/-^ mice. Interestingly, we noticed that cell proliferation genes (Klf4, Atf3, Grn) and their pathways including ERK1 and ERK2 cascade, MAPK signaling, cell growth, etc. were enriched in epsin-deficient mice **(Figure S3e and f)**, which contributed to reduced inflammation, promoted the cellular repair phenotypes^41,42^, and regulation of extracellular metabolism^43,44^. This observation suggests that epsin depleted mice exhibit a rewiring of cellular heterogeneity by inhibiting foam cell formation, inflammation mechanisms, and recapturing cellular functions.

### SCIG machine learning approach reveals the complete spectrum of cell identity genes in the atherosclerosis and epsin deletion mice

The marker genes derived from single-cell data analysis represent only a subset of cell identity genes (CIGs), potentially limiting their ability to fully capture the complexity of cell type identity regulation. These marker genes were defined based on differential expression analysis between individual cell types and others. Consequently, there is a possibility of overlooking well-known CIGs that play significant roles in regulating cell identity. To overcome this, we employed a novel machine-learning algorithm, SCIG to uncover the cell identity genes at the single-cell level. This algorithm integrates RNA expression profiles and genetic sequence features to identify the complete spectrum of CIGs within a given cell type. By applying SCIG, we successfully identified CIGs in each cell type of both ApoE^-/-^ and epsin-deficient littermate mice **(Figure 1e-g, S4a, S5a-c)**. To investigate the relationship between cell identity gene regulation in ApoE^-/-^ and epsin-deficient mice across various cell types, we performed Pearson correlation analysis using SICG-derived cell identity gene scores. Among the different VSMC subtypes, the highest correlation, 0.58, was observed in contractile VSMCs, suggesting that the cell identity profiles of ApoE^-/-^ and epsin-deficient mice are comparable in this subtype **(Figure 1f and 1g)**. Conversely, the lowest correlation, 0.38, was found in fibroblast-like VSMCs, indicating substantial alterations in cell identity profiles and functions due to epsin deficiency. Additionally, correlations of 0.29 and 0.33 were observed in macrophages and endothelial cells, respectively **(Figure 1f and 1g)**. These findings suggest that epsin significantly impacts the regulation of cell identity in these cell types, potentially playing a crucial role in the progression of atherosclerosis.

Pathway enrichment analysis revealed that the CIGs derived from SCIG exhibited a stronger association with their respective cell identities and functions **(Figure S4b-h and S5a-c)**. For example, CIGs identified by VSMC-like, and endothelial cells were enriched with smooth muscle cell proliferation, muscle contraction (**Figure S4b**), and endothelial cell development, angiogenesis (**Figure S5b**), respectively. Additionally, the identified cell identity genes in macrophages were enriched with cholesterol efflux, glucose metabolic process, foam cell differentiation and other macrophages related functional pathways (**Figure S5a**). These findings highlight the effectiveness of SCIG in identifying comprehensive sets of CIGs, providing deeper insights into cellular identity regulation in atherosclerosis context.

### Deletion of Epsins rescued the phenotypic switch between VSMC and Macrophage populations and EndMT

We observed significant changes in cell population changes between atherosclerosis and epsin-deleted mice fed a Western diet. Specifically, VSMC, endothelial cells, and mesenchymal cells showed increased populations, while macrophage populations decreased upon epsin depletion (**Figure 1c**). In addition to this, we observed that the expression of macrophage marker genes is upregulated in VSMC cells, endothelial cells, and mesenchymal cells of the atherosclerosis condition, whereas they are all downregulated when epsin is depleted (**Figure 2b**). This indicates that the elevated expression signals of these macrophage marker genes play a role in altering other cell identity functions. Based on these observations, we are interested in investigating the cellular dynamics under these conditions, and we conducted RNA velocity analysis independently. Remarkably, we identified two cell-type transitions in atherosclerosis mice: i) VSMC cells, particularly modulating VSMC2, exhibited a stronger transition toward macrophages, and ii) Modulating VSMC1 cells associated with inflammatory responses, and it transitioned into fibroblast-like VSMC and mesenchymal cells. However, these cell-type transitions were significantly rescued by deletion of epsins (**Figure 2a**).

**Figure 2:**
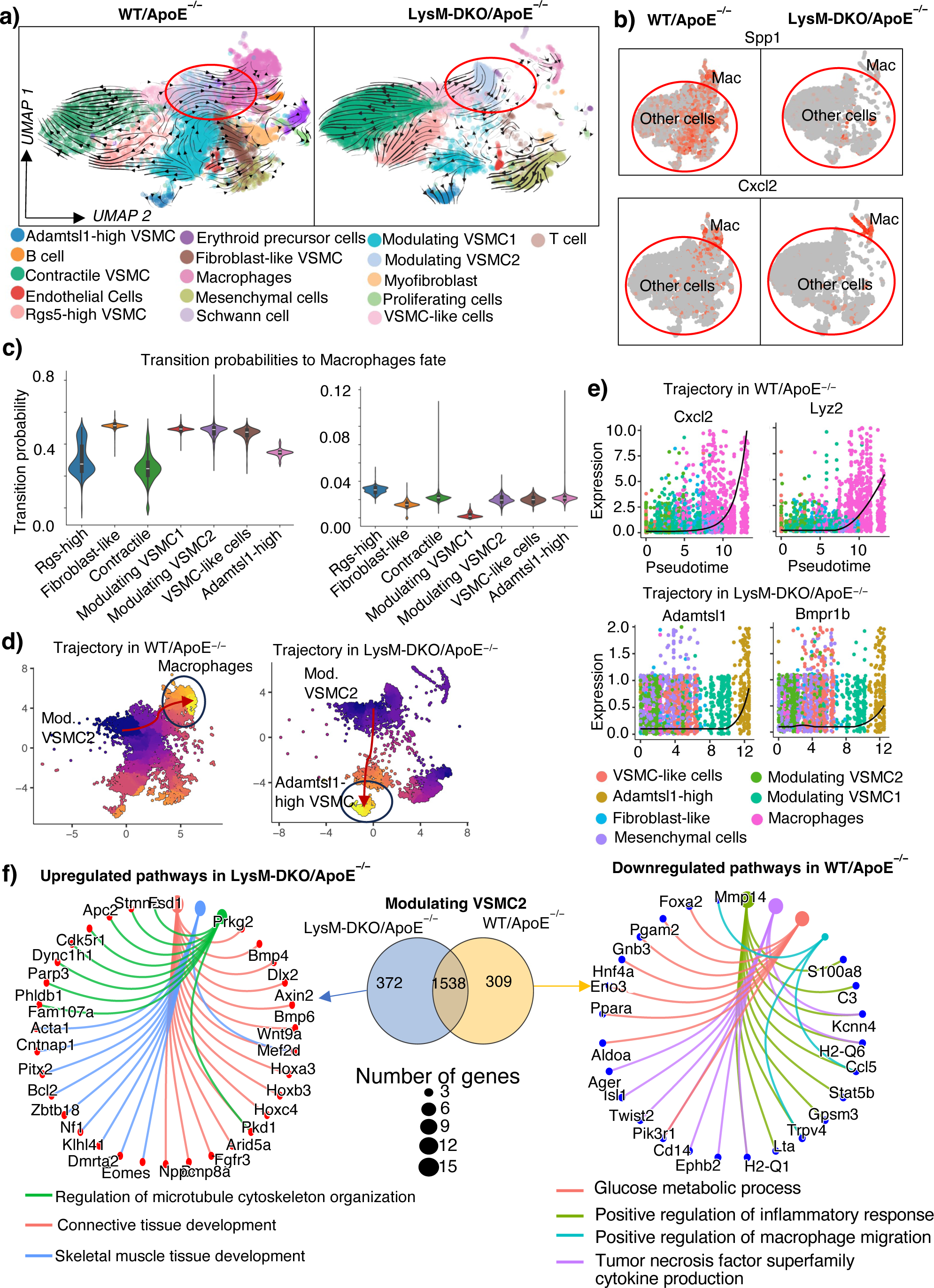
Deletion of macrophage-epsins reduces foam cell formation by inhibiting the crosstalk between VSMCs and macrophages. a) RNA velocity analysis reveals VSMCs transitioning to a macrophage-like phenotype in WT/ApoE^-/-^ mice, significantly reduced upon epsin deletion in LysM-DKO/ApoE^-/-^ mice. b) Expression levels of macrophage markers Spp1 and Cxcl2 in other cell types induce the VSMC to macrophage transition in WT/ApoE^-/-^, but not in the LysM-DKO/ApoE^-/-^ mice. c) Transition probabilities of various VSMC subtypes towards macrophages in both WT/ApoE^-/-^ and LysM-DKO/ApoE^-/-^ conditions. d) Monocle trajectory analysis confirming the VSMC-to-macrophage transition in WT/ApoE^-/-^ mice, fully inhibited in LysM-DKO/ApoE^-/-^ mice. e) Gene expression of macrophage and Adamtsl1-high VSMC marker genes upregulated in the trajectories of WT/ApoE^-/-^, and LysM-DKO/ApoE^-/-^, respectively. f) SCIG comparison identifying unique cell identity genes in modulating VSMC subtypes under each condition and their associated functional pathways.

Furthermore, we quantified the probability of each cell transition using CellRank and found that atherosclerosis mice have the maximum transition probability of 0.57 for modulating VSMC2 to macrophages, followed by modulating VSMC1, VSMC-like cells and fibroblast-like VSMC, all achieving probabilities of more than 0.5 (**Figure 2c left**). Conversely, these cell types did not exhibit a stronger transition towards macrophages when epsin was deleted (**Figure 2c right**). This suggests that epsin depletion primarily inhibits the transition of VSMC cells to macrophages.

### Monocle trajectory analysis also reveals the role of epsins in altering the ApoE^-/-^ mice cellular dynamics

We also conducted a monocle trajectory analysis to explore how changes in gene expression patterns influence cell phenotype switches between ApoE^-/-^ and epsin-deletion conditions. We selected modulating VSMC2 cells as root or start cells and their trajectory showed a transition toward macrophages in the ApoE^-/-^ condition (**Figure 2d left**). Additionally, we observed significant upregulation of macrophage marker genes such as Cxcl2, and Lyz2 in this trajectory (**Figure 2e top**). However, this cell type transition trajectory was disrupted when epsins were deleted and the modulating VSMC2 cells exhibited a trajectory towards Adamtsl1-high VSMC populations involved in extracellular matrix assembly functions (**Figure 2d right**). In case of the epsin-deleted condition, changes in gene expression, particularly for Adamtsl1 and Bmpr1b, led to the acquisition of modulating cells towards an Adamtsl1-high VSMC phenotype (**Figure 2e bottom**).

### SCIG revealed the condition-specific cell identity genes that maintain the VSMC identity

Our newly developed SCIG algorithm identified the complete list of cell identity genes in a given cell type. Leveraging this, we analyzed our SCIG-identified cell identity genes for Modulating VSMC2 cells in both ApoE^-/-^ and epsin-deleted condition. We observed that approximately 20% of cell identity genes were significantly different between wild-type and epsin-knockout conditions (**Figure 2f center**). Moreover, pathway enrichment analysis on these unique cell identity genes showed that skeletal muscle cell development, cytoskeleton organization, and connective tissue development were associated with epsin-deleted conditions (**Figure 2f left**), whereas in ApoE^-/-^ condition, macrophage migration, glucose metabolic process, inflammatory response, and cytokine production pathways were enriched (**Figure 2f right**). This observation was consistent with a pathway analysis of differentially expressed genes between two conditions (**Figure S6a)**. These findings indicate that epsin deletion strongly influences the maintenance of VSMC cells’ identity and functions.

### Deletion of macrophages epsin reverses the transition of endothelial cells to mesenchymal cells

Endothelial-to-mesenchymal transition (EndMT) is a process wherein endothelial cells lose their identity and differentiate into mesenchymal cells, leading to the rapid formation of unstable atherosclerotic plaques ^45,46^. Our cellular dynamics analysis, in line with previous findings, revealed that endothelial cells are more prone to transitioning into mesenchymal cells, and fibroblast-like VSMC cells when epsins are present **(Figure 3a, left)**. However, this cellular phenotype switching was reversed in epsin-deficient mice **(Figure 3a, right)**. Further, monocle trajectory analysis captured the gene expression changes, showing downregulation of endothelial cell markers and upregulation of mesenchymal cell marker genes, thereby affirming the EndMT process is facilitated by epsins in Western diet-fed atherosclerosis mice **(Figure 3b left and c left)**. Conversely, the gene expression profiles of EndMT process were reversed to a mesenchymal-to-endothelial transition (MendT) phenotype in epsin-deficient mice **(Figure 3b right and c right)**.

**Figure 3:**
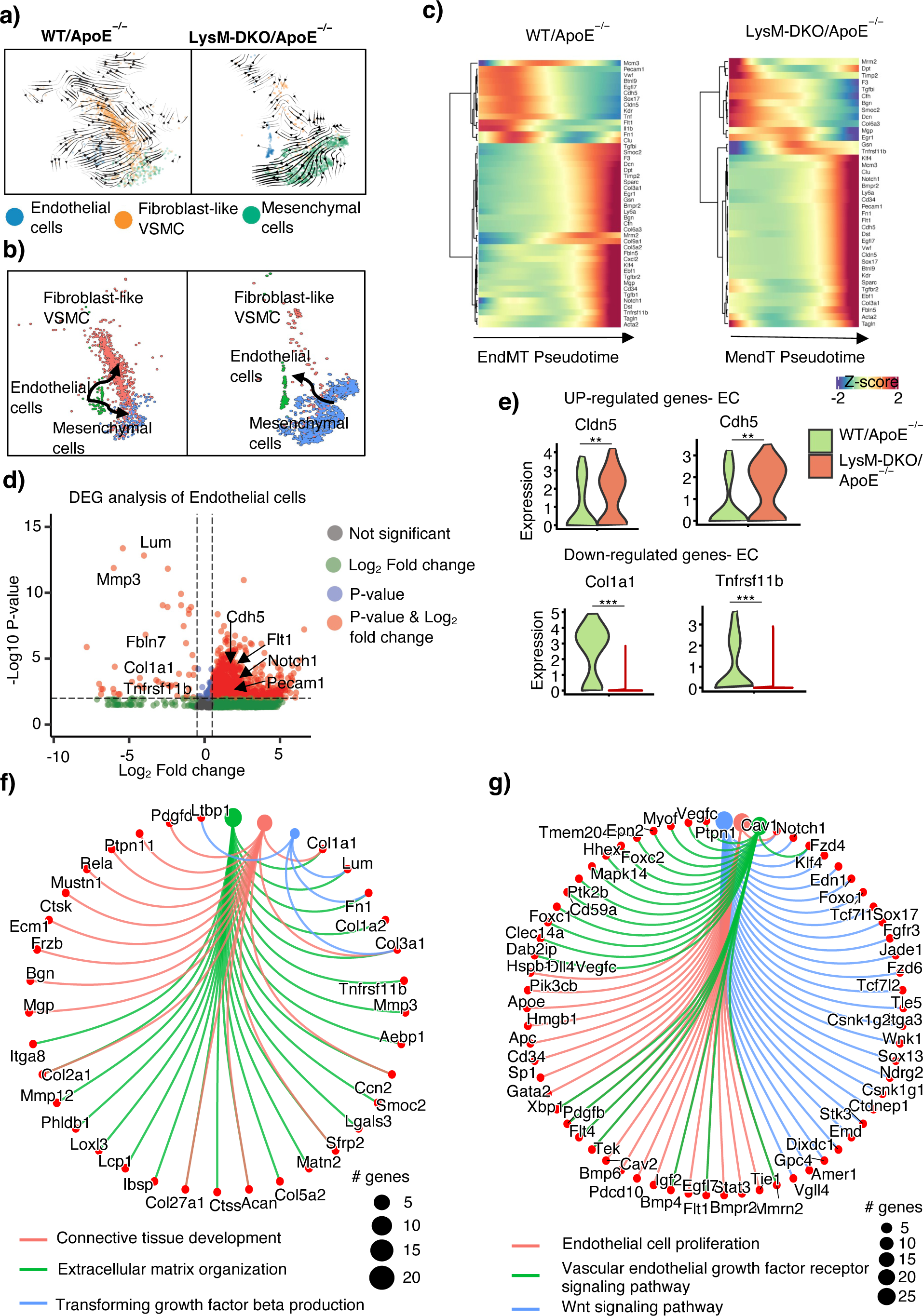
Endothelial to mesenchymal cell transition diminished and reversed by epsins deletion. a) RNA velocity analysis reveals that epsin promotes EndMT in WT/ApoE^-/-^ mice, whereas its deletion facilitates a transition from mesenchymal to endothelial cells. b) Monocle3 trajectory analysis confirms these cell transitions observed via RNA velocity. c) Heatmap demonstrating downregulation of endothelial cell markers in WT/ApoE^-/-^ mice and upregulation in LysM-DKO/ApoE^-/-^ mice, corroborating the EndMT and MendoT processes, respectively. d) Volcano plot highlighting differentially expressed genes in endothelial cells between WT/ApoE^-/-^ and LysM-DKO/ApoE^-/-^ mice. e) Violin plot illustrating the upregulation of endothelial cell markers (Cdh5, Cldn5) in LysM-DKO/ApoE^-/-^ mice and mesenchymal markers (Col1a1, Tnfrsf11b) in WT/ApoE^-/-^ mice. f) Functional pathway analysis showing genes upregulated in WT/ApoE^-/-^ mice associated with mesenchymal and smooth muscle development. g) Genes upregulated in LysM-DKO/ApoE^-/-^ mice are enriched in endothelial cell functional pathways. *Note: P values determined by the Wilcoxon test. *, p-value < 0.05; **, p-value < 0.01; ***, p-value < 0.001*.

Furthermore, we identified differentially expressed genes in endothelial cells under two conditions and observed that the gene expression patterns of pan-endothelial cell markers (Flt1, Pecam1, Cdh5, Cldn5) were significantly upregulated in the epsin-knockout condition compared to the ApoE^-/-^ condition **(Figure 3d, e and S2d)**. Additionally, we analyzed the SCIG-derived cell identity genes of endothelial cells and found that 602 and 363 identity genes were uniquely identified in wild-type and epsin-deletion conditions, respectively **(Figure S6b)**. Pathway enrichment analysis revealed that the unique endothelial cell identity genes in the epsin-deletion condition were enriched for VEGF receptor pathways, endothelial proliferation, endothelium development, angiogenesis, and WNT signaling **(Figure 3g and S6b)**. In contrast, the unique identity genes in ApoE^-/-^ mice were associated with connective tissue development, extracellular matrix organization, transforming growth factor beta production, and fibroblast growth factor receptor signaling pathways **(Figure 3f and S6b).** These findings reaffirm that an atherosclerosis condition alters the normal functions of endothelial cells and promotes the EndMT process, which is reversed when epsin is deleted.

### Macrophage-specific epsin promotes the inflammation in fibroblast-like VSMC cells

Through differential expression analysis of fibroblast cells, we identified elevated gene expression profiles related to inflammation and cytokine production (Tgfbr3, Ccl19) and repression of fibroblast functional genes (Ccn1, Mef2c) in WT/ApoE^-/-^ mice **(Figure S6c)**. Additionally, exploration of cell identity genes in the fibroblast population also revealed that the WT/ApoE^-/-^ condition-specific cell identity genes are often involved in autophagy, inflammatory responses, and cytokine responses **(Figure S6d)**. Conversely, the unique cell identity genes in the epsin-deletion condition were associated with fibroblast proliferation, activation, and muscle cell development pathways **(Figure S6d)**. These findings suggest that epsin plays a predominant role in inducing inflammation in fibroblasts, thereby affecting their cellular identity and functions.

### Cell-cell communications reveal that epsin deletion inhibits the inflammatory and cytokine signaling

We utilized NicheNet to explore the ligand-target genes mediated signals between macrophages and VSMC cells. Specifically, we defined “macrophages” and “VSMC subtypes” as sender and receiver cells, respectively. In WT/ApoE^-/-^ mice, ligands including Lgals3, II1b, C1qa, and C1qb were identified as potential ligands in macrophages **(Figure 4a left)**, known to be associated with inflammatory response and cytokine production. These ligands regulate transcription profiles of 49 genes such as Cxcl2, Tnfrsf11b, Ptgs2, and Icam1 in receiver VSMC cells **(Figure 4b and c left)**. The identified ligand-target gene mediated signals were mainly associated with inflammatory response, cytokine production, and regulation of myeloid differentiation. Conversely, in epsin-depleted conditions, ligands such as Abca1, Grn, and Psap were identified as top ligands in macrophages **(Figure 4a right)**, associated with clearing cholesterol and fat from the cells^47^. These ligands regulate 45 target genes (Aff4, Ccn1, Egr1) in VSMC cells, enriching ligand-target genes communication signals involved in removing cholesterol from the cells and regulating VSMC’s normal physiological functions **(Figure 4c right)**. This result indicates that inflammatory and cytokine mediated signals are significantly downregulated by the deletion of macrophage epsins.

**Figure 4:**
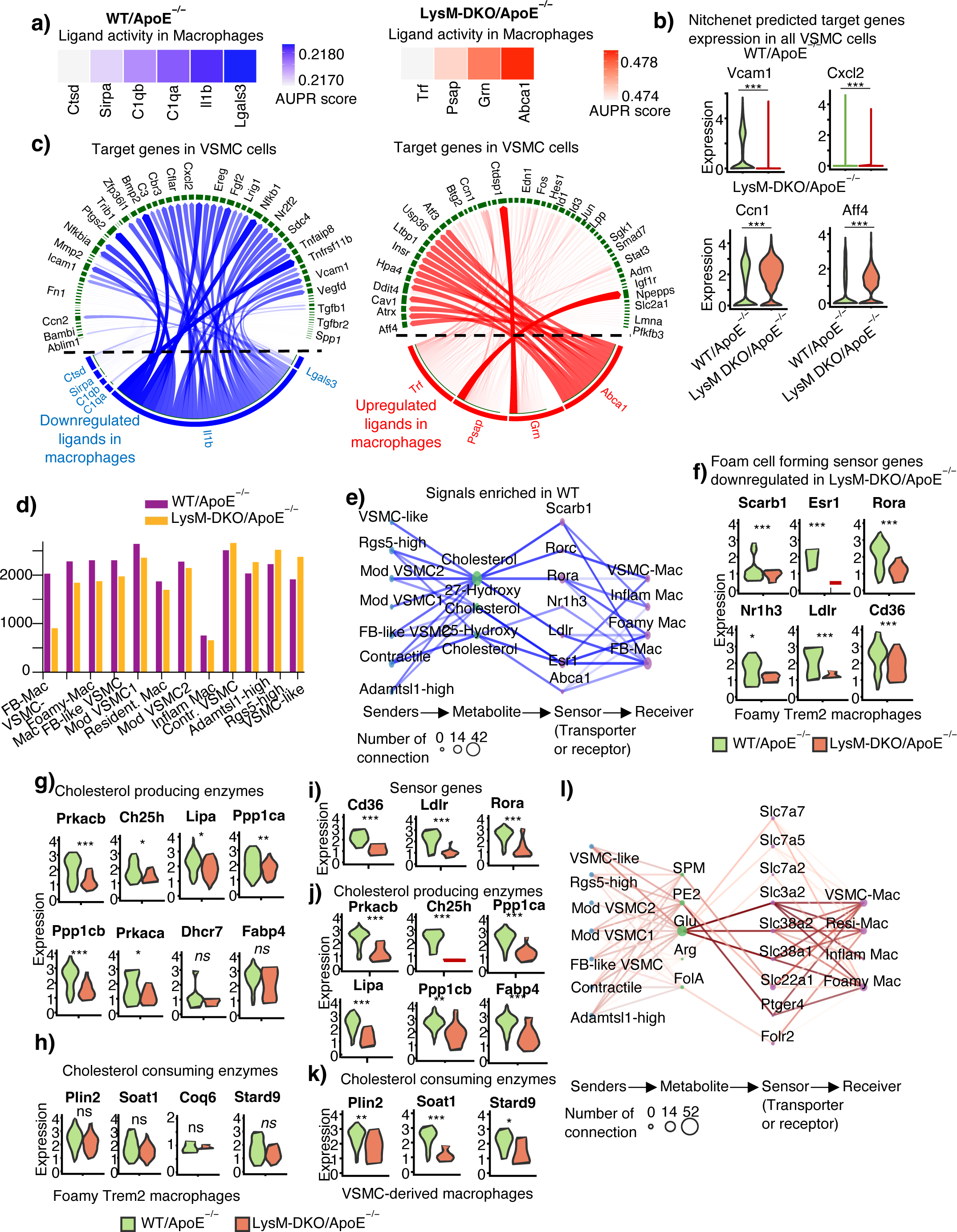
Exploring the molecular mechanisms of foam cell formation through the protein-ligand and metabolite-mediated cell-cell communications. a) NicheNet analysis identifies potential macrophage ligands that modulate inflammatory signaling in WT/ApoE^-/-^ mice and reduce foam cell formation in LysM-DKO/ApoE^-/-^ mice. b) Expression of predicted target genes in VSMC cells for these identified ligands shown in the violin plot. c) Circos plot illustrating the down-regulated and upregulated cell-cell communication signals in epsin-deficient mice. d) MEBOCOST analysis reveals metabolite-mediated cell communication signals in both WT/ApoE^-/-^ and LysM-DKO/ApoE^-/-^ mice. e) Predominant cell-cell communication patterns observed in WT/ApoE^-/-^ mice. g) Enriched expression of metabolite-producing enzymes in WT/ApoE^-/-^ mice. f-h) Expression level of cholesterol sensors, producing enzymes, consuming enzymes in WT/ApoE^-/-^ and LysM-DKO/ApoE^-/-^ mice of foam cells. i-k) Expression level of cholesterol sensors, producing enzymes, consuming enzymes in WT/ApoE^-/-^ and LysM-DKO/ApoE^-/-^ mice of VSMC-derived macropahge cells. l) Predominant cell-cell communication patterns observed in LysM-DKO/ApoE^-/-^ mice. *Note: P values determined by the Wilcoxon test. *, p-value < 0.05; **, p-value < 0.01; ***, p-value < 0.001*.

Through RNA velocity and trajectory analysis, we observed significant crosstalk between endothelial cells and mesenchymal, fibroblast-like VSMC **(Figure 3a, b)**. Specifically, the EndMT and MendT processes were observed in WT/ApoE^-/-^ and LysM-DKO/ApoE^-/-^ mice, respectively. We performed cell-cell communication analysis between these cell types to explore the underlying molecular mechanisms. Interestingly, we found that ligands such as Tgfb1, Tgfb2, and Bmp4 from mesenchymal and fibroblast-like VSMCs activate several EndMT driver genes, including Fn1, Col1a1, Tagln, Spp1, and Tnfrsf11b, in endothelial cells of WT/ApoE^-/-^ mice **(Figure S7a)**. These target genes exhibited higher expression levels in WT/ApoE^-/-^ mice compared to epsin-deficient mice **(Figure S7c)**. Remarkably, the EndMT process was reversed by deleting macrophage epsins. We observed that ligands such as Jam2, Mmrn2, Dll1, and Dll4 in endothelial cells induce the MendT process by targeting the Fos, Klf4, Jun, Id3, and Nrp1 genes **(Figure S7b)**. Notably, the expression of these genes is upregulated in LysM-DKO/ApoE^-/-^ mice **(Figure S7d)**.

### MEBOCOST identifies that macrophage-specific epsins reduce cholesterol and glucose-mediated cell-cell communications

We utilized our MEBOCOST^39^ algorithm to identify potential metabolite-mediated cell communication signals at the single-cell level. During this computation, MEBOCOST uses RNA expression signatures of metabolite-producing and consuming enzymes as well as sensors (transporters/receptors) expression. Applying MEBOCOST to our datasets, we observed a reduction in the number of metabolite-mediated cell signals in modulating VSMC1, VSMC2, and fibroblast-like VSMC cells, whereas other VSMC subtypes exhibited a higher number of communications in epsin-deficient mice (**Figure 4d)**. Notably, among all VSMC subtypes, modulating VSMC1, VSMC2, and fibroblast-like VSMC cells were associated with inflammation, migration, cytokine production, and the phenotypic switch of VSMCs into macrophages. The reduced number of cell communications in these cell types indicates that epsin deletion may inhibit these processes and help maintain cellular identity.

To validate this hypothesis, we conducted a differential communication analysis between WT/ApoE^-/-^ and LysM-DKO/ApoE^-/-^ mice. Our analysis revealed that cholesterol^48,49^, and its derivatives (27-hydroxycholesterol^50^ and 25-hydroxycholesterol^51,52^) mediated cell signals were associated with WT/ApoE^-/-^ mice (**Figure 4e)** but not with epsin-deficient ones (**Figure 4i)**. These results suggest that the downregulation of cholesterol-mediated cell signals significantly reduces foam cell formation, consistent with our findings in normal diet mice^28^. Additionally, uridine diphosphate glucose^53,54^, sphingosine-1-phosphate^55^, hypoxanthine ^56,57^, fructose^58,59^, and adenosine monophosphate^60^ signals were also implicated in regulating foam cell formation, promoting atherosclerosis progression (**Figure S8a)**. The expression of cholesterol-producing and consuming enzymes, as well as their sensors, was downregulated in various VSMC and macrophage subtypes (**Figure 4f-k and Figure S8b-d)**. For example, the expression levels of cholesterol-transporting genes such as Cd36, Ldlr, and Rora (**Figure 4f and i**), cholesterol-producing enzymes such as Prkacb, Lipa, and Ch25h (**Figure 4g and j**), and cholesterol-consuming enzymes Plin2, Soat1, and Stard9 (**Figure 4k**) were significantly reduced in foamy and VSMC-derived macrophages of LysM-DKO/ApoE^-/-^ mice. In the case of epsin-deficient mice, prostaglandin E2^61^, L-Glutamine^62^, L-Arginine^63,64^, folic acid^65^, and spermine^66–68^ mediated signals were upregulated (**Figure 4l)** and these signals have been reported to rescue foam cell formation and plaque instability, ultimately delaying atherosclerosis progression.

## Discussion

Atherosclerosis is a chronic inflammatory condition marked by the accumulation of lipids and fat molecules (plaques formation) within the arterial walls. This pathological buildup leads to significant vascular dysfunction, contributing to the development of coronary heart disease, ischemic stroke, and peripheral artery disease^69,70^. Unhealthy dietary patterns are a major risk factor for the onset and progression of atherosclerosis^71^. Consequently, identifying novel therapeutic targets and developing effective intervention strategies are essential for mitigating the impact of this disease.

Recently, we demonstrated that epsin 1 and 2 in macrophages play pivotal roles in cholesterol transport, promoting foam cell formation, which is positively correlated with atherosclerosis progression^27,28,30^. To understand the functional impact of macrophage epsins in atherosclerosis progression, we constructed two mice models fed a Western diet for 16 weeks: one mimicking atherosclerosis progression (WT/ApoE^-/-^) and the other inhibiting this progression by targeting macrophage epsins (LysM-DKO/ApoE^-/-^). Next, we analyzed the transcriptomic profiles of these cells using single-cell RNA sequencing technology. Our bioinformatic data analysis revealed significant cellular heterogeneity present in vascular smooth muscle cells (VSMCs) and macrophages, with functional enrichment analysis indicating distinct cellular functions. Additionally, we employed our newly developed algorithm, SCIG, to decipher the cell identity profiles of each cell type, providing deeper insights into cell identity regulation. Notably, WT/ApoE^-/-^ mice, modulating VSMC1 cells were involved in inflammation and migration, while modulating VSMC2 cells predominantly underwent phenotypic switches into macrophages. These findings suggest that modulating VSMCs is a major source of foam cell formation and inflammation thereby accelerating disease progression. Conversely, LysM-DKO/ApoE^-/-^ mice showed increased populations of VSMC-like cells, contractile cells, Adamtsl1-high cells, Rgs-high cells, and resident-like macrophages while decreasing populations of modulating VSMC cells, VSMC-derived macrophages, fibroblast-derived macrophages, foamy macrophages, and inflammatory macrophages. Furthermore, RNA velocity analysis indicated that the regulatory role of epsins in the VSMC-to-macrophage transition was diminished. These results suggest that deleting epsins in macrophages significantly enhances the maintenance of VSMC identity and function while inhibiting inflammatory cell signals.

The endothelial-to-mesenchymal cell transition contributes to foam cell formation and inflammatory signaling. Our atherosclerotic mice model exhibited prevalent EndMT phenotypic changes, as confirmed by RNA velocity and gene expression changes in the trajectory analysis. We observed a significant downregulation of known endothelial cell markers and upregulation of mesenchymal cell markers in WT/ApoE^-/-^ mice. In contrast, the gene expression patterns indicating EndMT were completely reversed to mesenchymal-to-endothelial transition in epsin-deficient mice. This confirms that macrophage epsins play a critical role in endothelial and mesenchymal cell functions.

Cell-cell communication analysis revealed that inflammatory signals were primarily driven by ligands (Il1b, C1qa, Lgals3) in macrophages, which altered the functions and identities of VSMCs and other cell types. However, these signals were downregulated and rewired in epsin-deficient mice. Additionally, Abca1, Psap, and Grn ligand-mediated cell communications were upregulated in LysM-DKO/ApoE^-/-^ mice, reducing inflammation and maintaining lipid homeostasis. Our metabolite-mediated cell communication analysis showed that cholesterol, glucose, and hypoxanthine-mediated signals were associated with WT/ApoE^-/-^ mice, enhancing foam cell growth. Conversely, deleting macrophage epsins reduced these signals and promoted prostaglandin E2, L-glutamine, and folic acid-mediated communications, delaying atherosclerosis progression.

In this study, we revealed the regulatory role of macrophage epsins and their molecular mechanisms in foam cell formation, which are ultimately associated with atherosclerotic lesion development. Our strategy of targeting epsins in macrophages in atherosclerosis encompasses several approaches to inhibit foam cell growth: i) increasing the population of VSMCs while decreasing foam cells and inflammatory macrophages, ii) maintaining the cellular identity and functions of VSMCs, endothelial cells, and fibroblasts by preventing abnormal cell transitions, iii) downregulating inflammation and cytokine production-mediated cell signals, and iv) reducing cholesterol and glucose-mediated cell communications. In summary, we found that inhibiting macrophage epsins serves as a predominant target for developing potential treatment strategies against various heart diseases, including ischemic stroke and coronary heart disease.

## Data Availability

The authors declare that all supporting data are available within the article and its Supplemental Material. Additional methods or data related to this study are available from the corresponding authors upon reasonable request. The single-cell RNA sequencing data generated in this study were deposited into the GEO database with the accession number GSE214414.

## Acknowledgments

This project is supported by R01HL146134 (H.C.) and American Heart Association Established Investigator Award and Transformational Project Award (H.C.), and in part by NIH grants R01HL137229, R01HL141853, R01HL156362, R01HL158097, R01HL162367, (H.C.), and R01GM125632 (K.F.C.)

## Author Contributions

H.C. and K.F.C. conceived the project. K.A. performed the bioinformatic data analysis and interpreted the data. K.C. and A.V. conducted the experimental verification and analysis. All the analysis was done under the supervision of H.C. and K.F.C. K.A. and K.C. wrote the manuscript with comments from H.C. X.G. and K.F.C. All authors reviewed and approved the manuscript.

## Supplementary Figure Legends

**Figure S1: Identification of major cell clusters and cell type annotations via single-cell RNA sequencing.**

a) Violin plot showing the stringent conditions used to filter out low-quality cells before downstream analysis. b) UMAP plot displaying the spatial distribution of identified major cell types in WT/ApoE^-/-^ and LysM-DKO/ApoE^-/-^ mice. c) Expression of known marker genes used for annotating the cell types. d) Comparison of cell population sizes between WT/ApoE^-/-^ and LysM-DKO/ApoE^-/-^ mice. e) qRT-PCR was performed to profile the expression of TNF-α gene (n=3), data are presented as mean±SD. Student’s t-test were used to analyze the data. *\*\*, p-value < 0.01; ***, p-value < 0.001*.

**Figure S2: Marker genes expression of WT/ApoE^-/-^ and LysM-DKO/ApoE^-/-^ mice.**

a-d) Aortas from WT/ApoE^-/-^ and LysM-DKO/ApoE^-/-^ mice were isolated, qRT-PCR was performed to check the expression of marker genes (n=3), data from a through e are presented as mean±SD. Student’s t-test were used to analyze the data. *\*\*\*, p-value < 0.001*.

**Figure S3: Sub-clustering analysis of macrophages reveals five major sub-cell types.**

a) UMAP plot showing the spatial positions of the five macrophage sub-types. b) Violin plot depicting the marker gene expression of each macrophage sub-type. c) qRT-PCR was performed to check the expression of marker genes (n=3). data are presented as mean±SD. Student’s t-test were used to analyze the data. *\*\*, p-value < 0.01; ***, p-value < 0.001*. d) Comparison of cell population sizes of each sub-type between atherosclerosis and epsin-deficient mice. e) Upregulation of cell proliferation pathways in resident-like macrophages of epsin-deficient mice. f) Expression levels of cell proliferation markers in the resident-like macrophage population.

**Figure S4: SCIG uncovered the complete spectrum of cell identity genes of VSMC subtypes and their functional annotations.**

a) Venn diagram showing the comparison of identified SCIG-derived cell identity genes between WT/ApoE^-/-^ and LysM-DKO/ApoE^-/-^. b-h) Bar plots comparing pathway enrichment analysis of SCIG genes shows the cell identity of VSMC subtypes, including VSMC-like cells (b), contractile cells (c), Rgs5-high cells (d), Adamtsl1-high cells (e), Modulating1 cells (f), Modulating2 cells (g), and fibroblast-like VSMCs (h).

**Figure S5: SCIG uncovered the cell identity gene profiles of macrophages, endothelial cells, and mesenchymal cells and their functional annotations.**

a-c) Bar plots comparing pathway enrichment analysis on identified SCIG-derived cell identity genes between WT/ApoE^-/-^ and LysM-DKO/ApoE^-/-^ for macrophages (a), endothelial cells (b), and mesenchymal cells (c).

**Figure S6: Pathway enrichment analysis of condition-specific cell identity genes (WT/ApoE^-/-^ or LysM-DKO/ApoE^-/-^ mice) revealed by SICG algorithm.**

a-b) Pathways associated with SCIG-derived specific cell identity genes in epsin-deficient and atherosclerotic mice for Modulating VSMC 2 (a) and endothelial (b) cells. c) Expression levels of both upregulated and downregulated genes in fibroblast-like VSMC cells of epsin-deficient mice. d) Bar plot showing pathway associations of differentially expressed genes in fibroblast cells. *Note: P values determined by the Wilcoxon test. *, p-value < 0.05; **, p-value < 0.01; ***, p-value < 0.001.*

**Figure S7: Nitchenet analysis identified potential ligand-target gene cell communications in EndMT and MendT process.**

a) Circos plot depicting ligand-target gene interactions in the EndoMT process. b) Circos plot depicting ligand-target gene interactions in the MendoT process. c) Violin plot showing the expression levels of target genes in EndoMT. d) Violin plot showing the expression levels of target genes in MendoT. *Note: P values determined by the Wilcoxon test. *, p-value < 0.05; **, p-value < 0.01; ***, p-value < 0.001.*

**Figure S8: MEBOCOST identified the glucose and other metabolites mediated cell communications associated with growth of foam cells.**

a) Glucose and other metabolites mediated signals predominantly observed in WT/ApoE^-/-^ mice. g) Enriched expression of metabolite-producing enzymes in WT/ApoE^-/-^ mice. b-d) Expression level of glucose and other metabolites sensors, in WT/ApoE^-/-^ and LysM-DKO/ApoE^-/-^ mice of foam cells and VSMC-derived macropahge cells. *Note:UDP: Uridine diphosphate glucose, S1P:Sphingosine 1-phosphate, AMP: Adenosine monophosphate, D-Fruc: D-Fructose, Hyp:Hypoxanthine. SPM: PE2: Prostaglandin E2, Glu: L-Glutamine, Arg: L-Arginine, FolA: Folic acid. P values determined by the Wilcoxon test. *, p-value < 0.05; **, p-value < 0.01; ***, p-value < 0.001.*

