## Supplementary figures and images for "Single-Cell Analysis Reveals Critical Role of Macrophage Epsin in Regulating Origin of Foam Cell in Atherosclerosis"

### Supplemental Figures

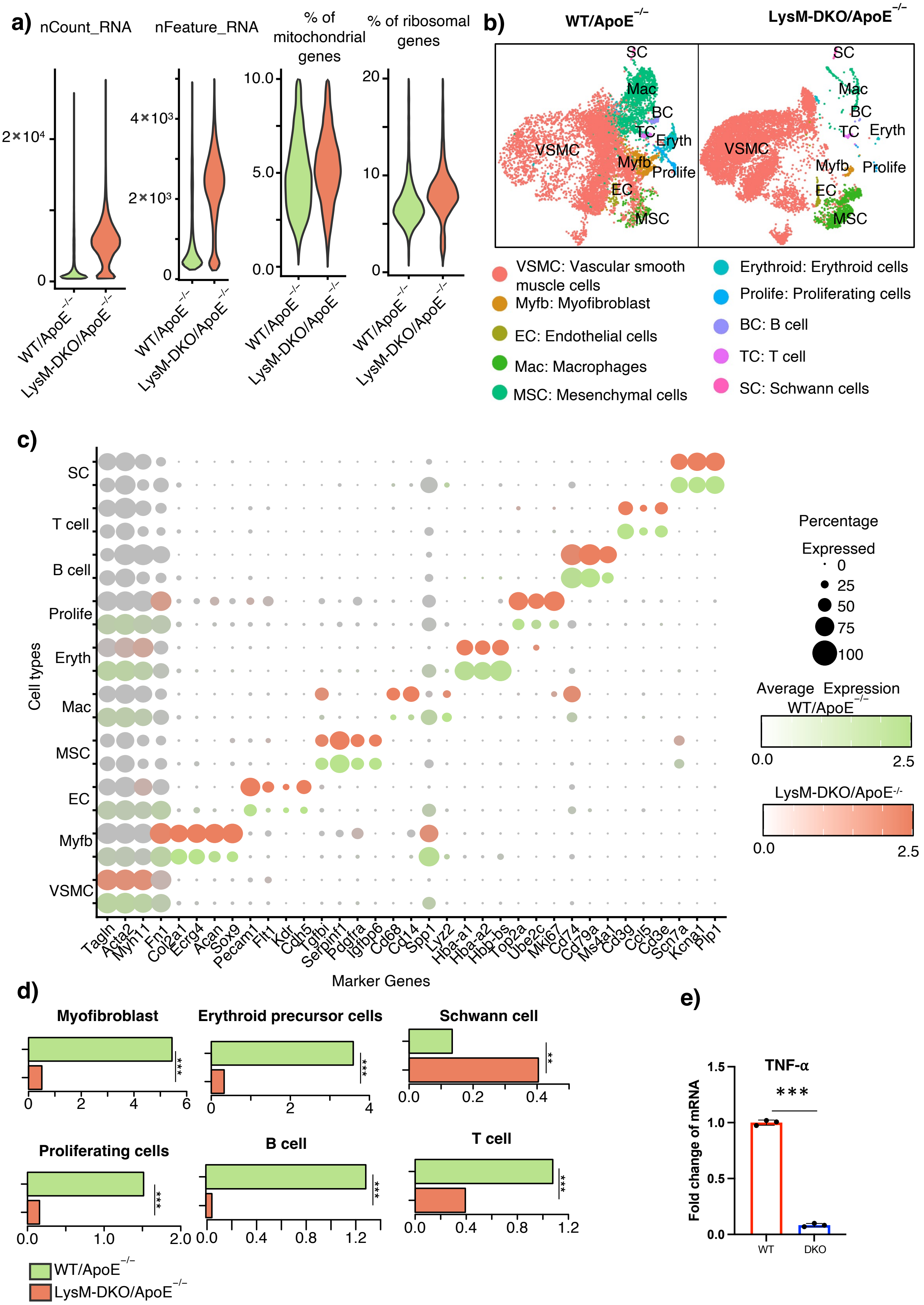

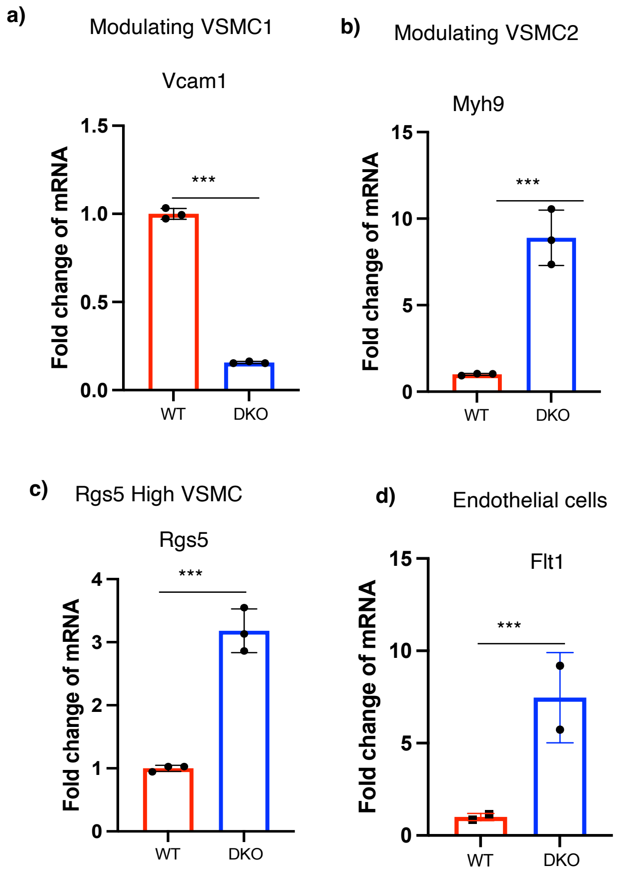

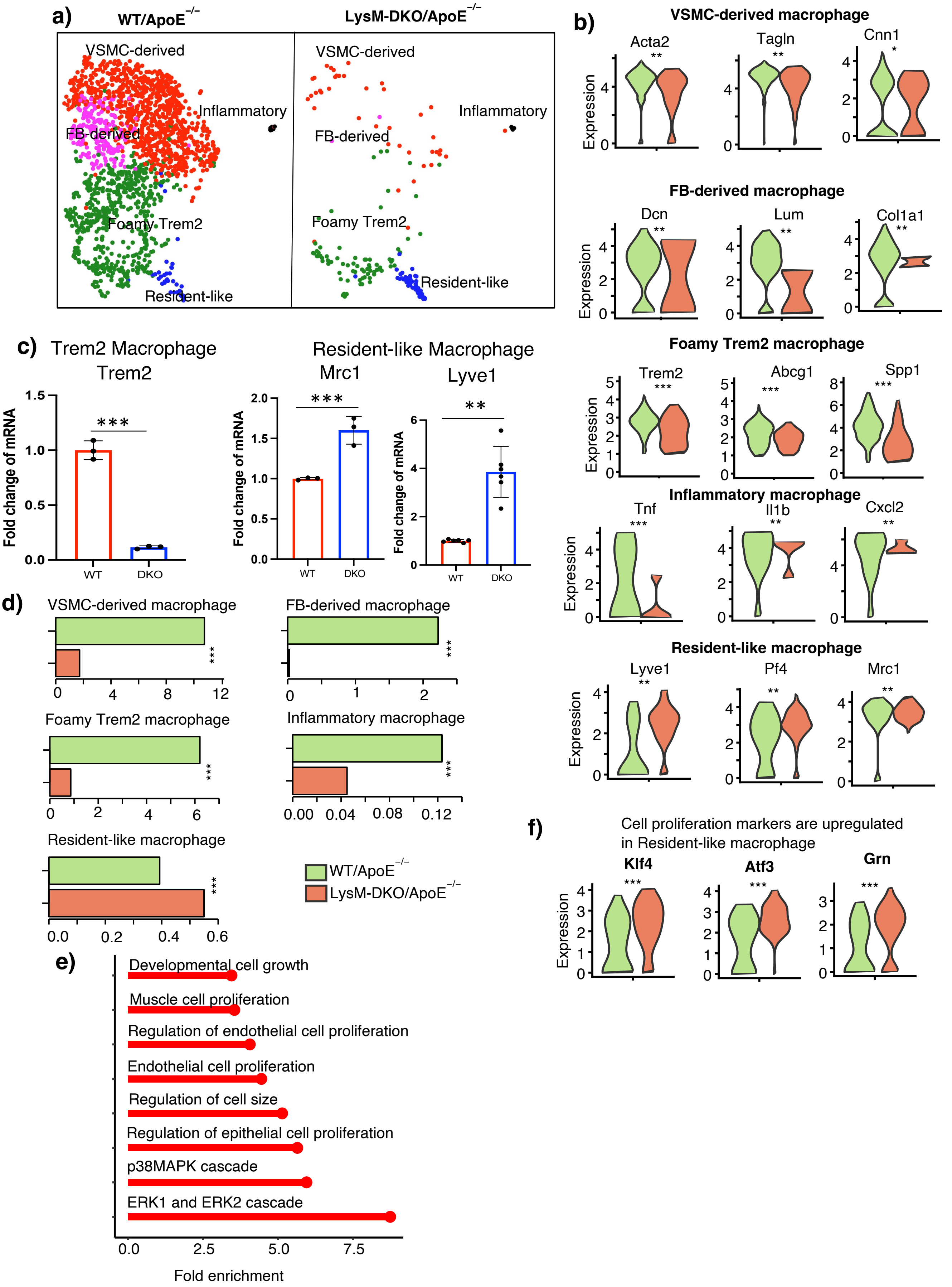

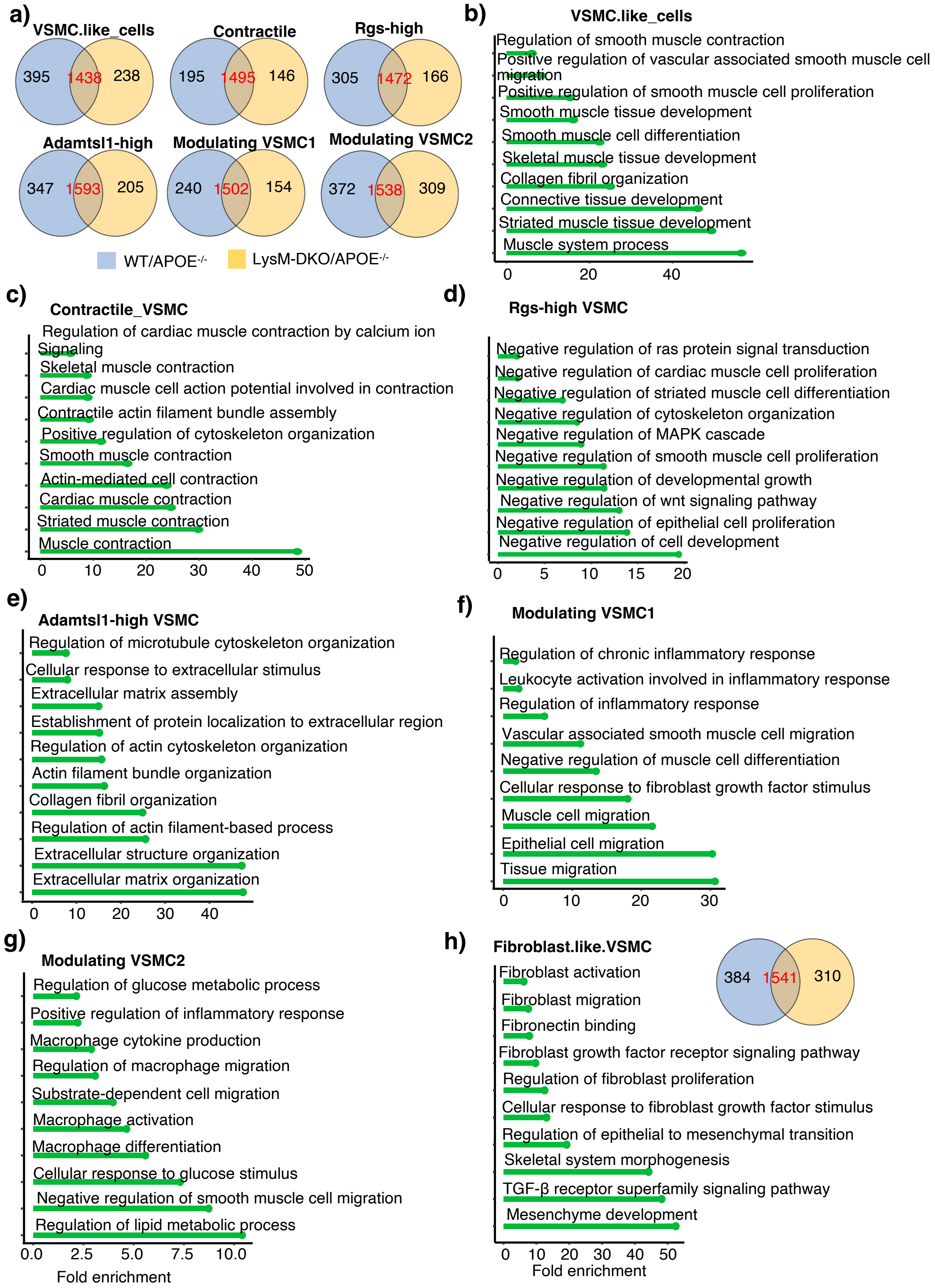

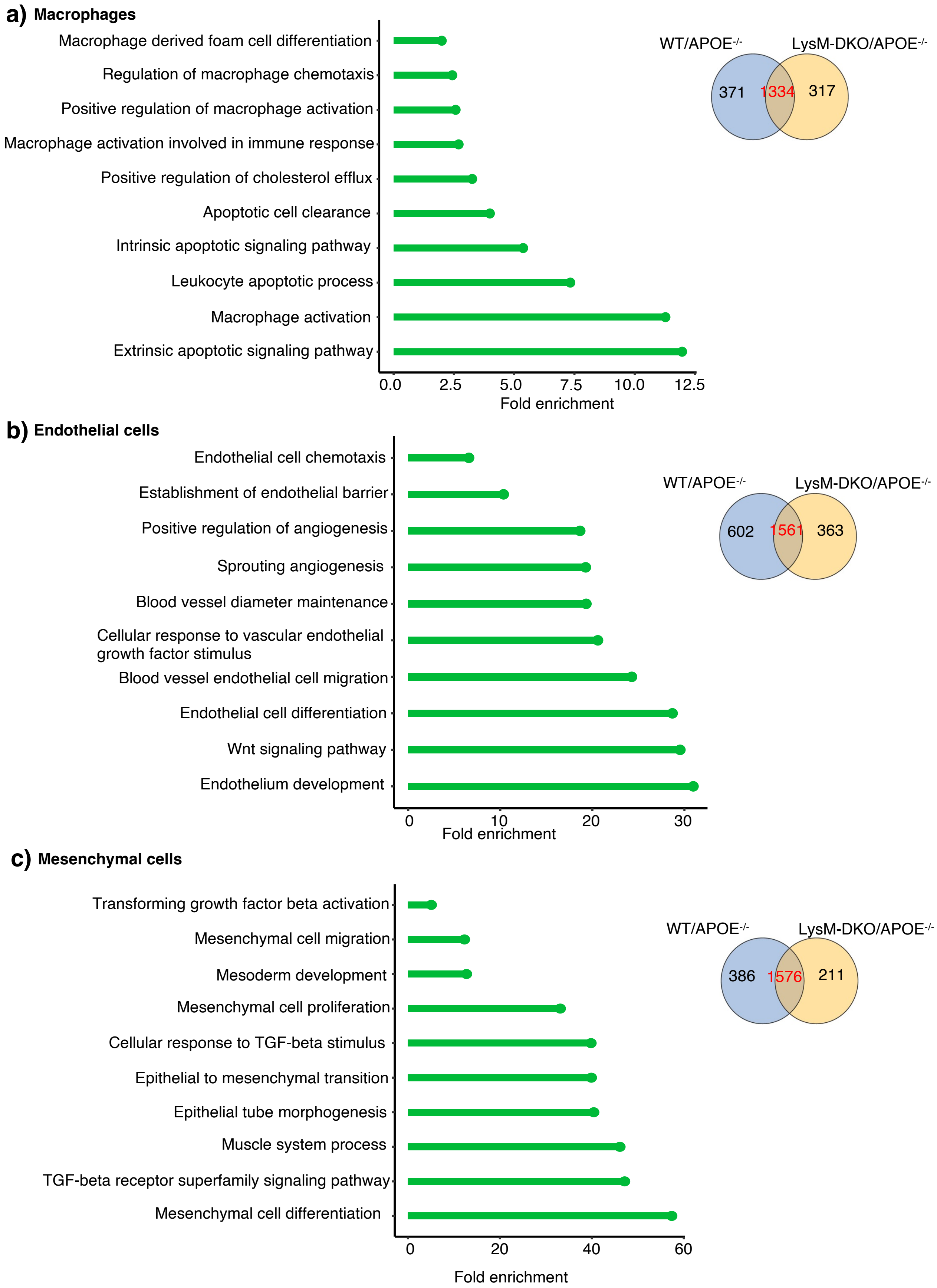

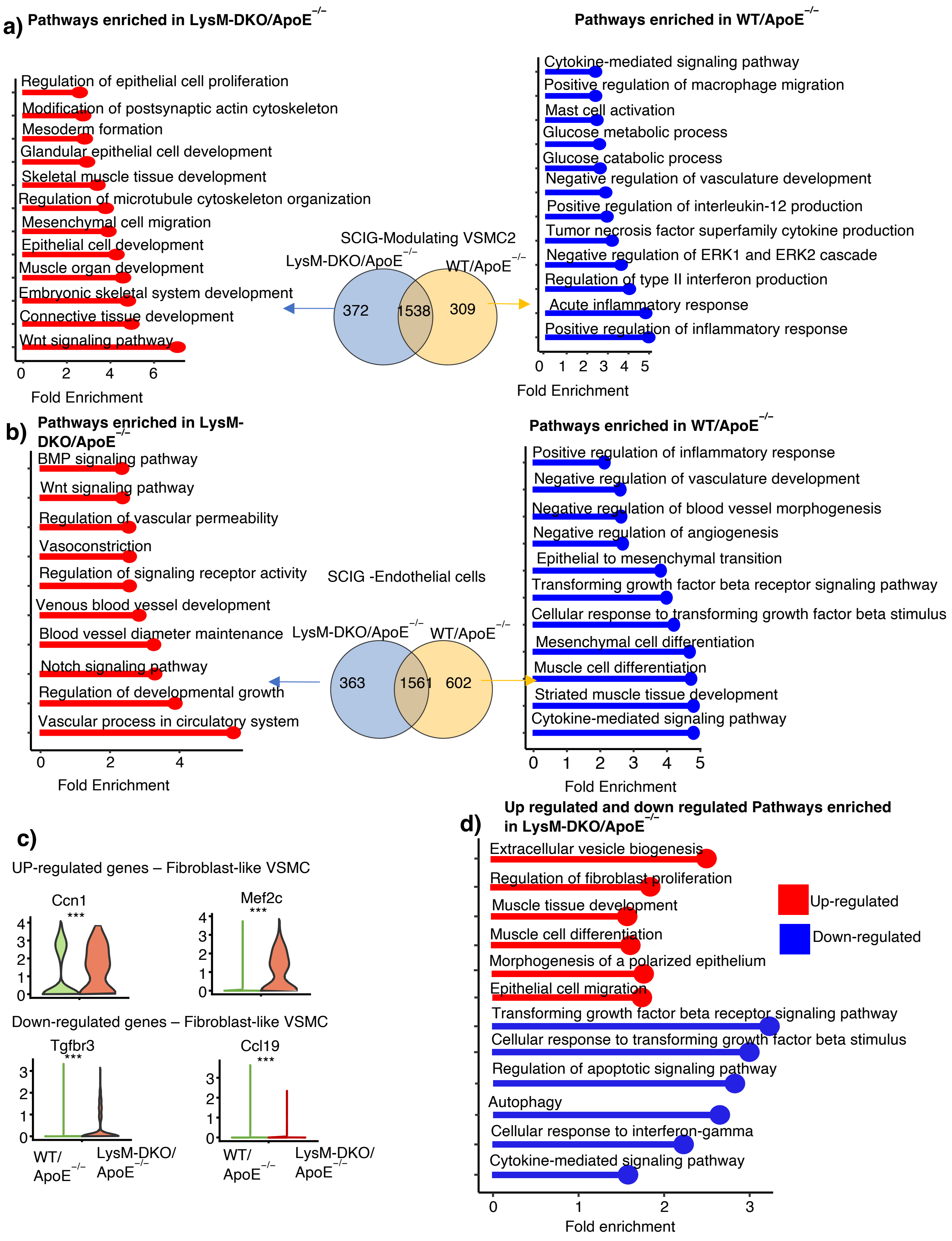

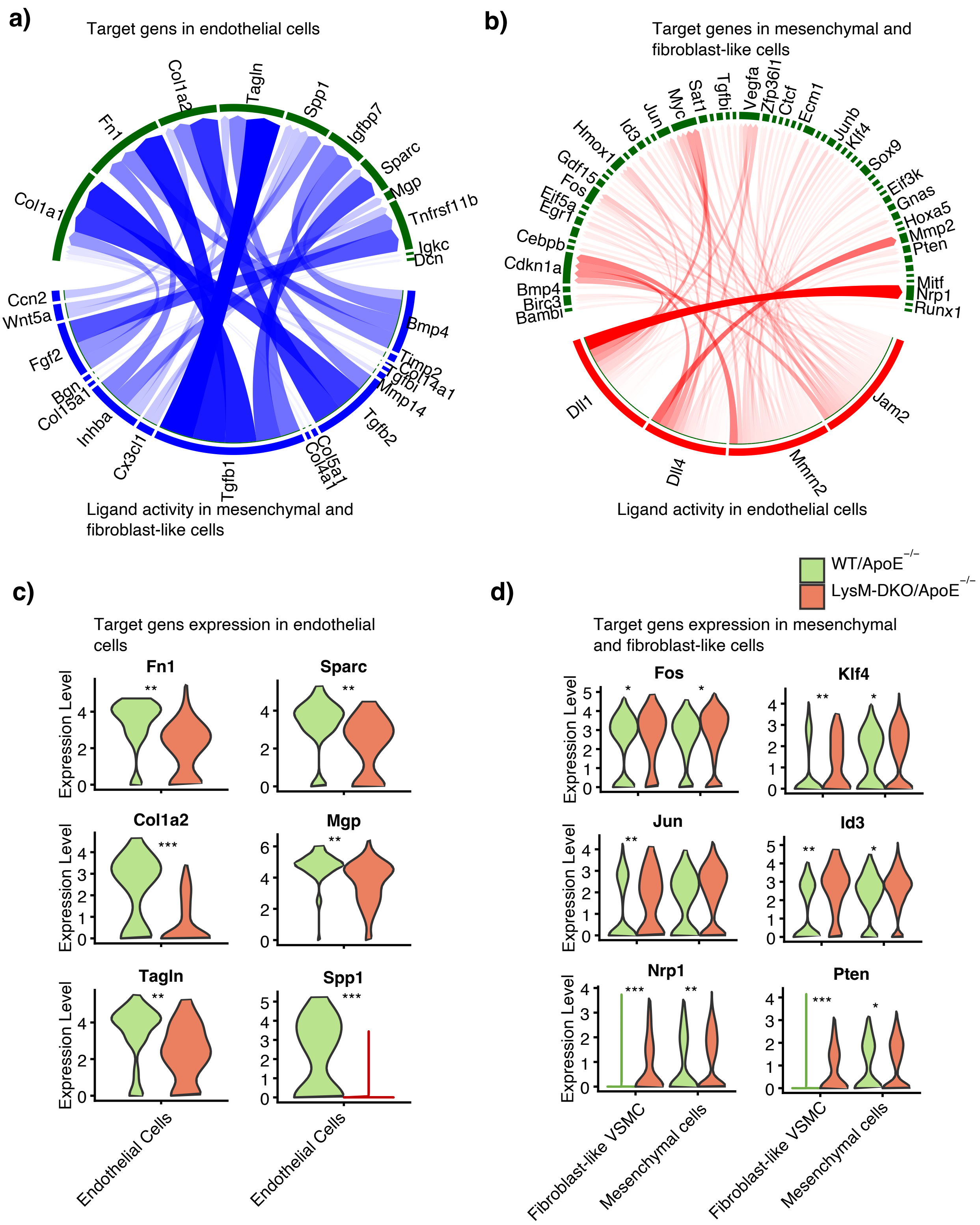

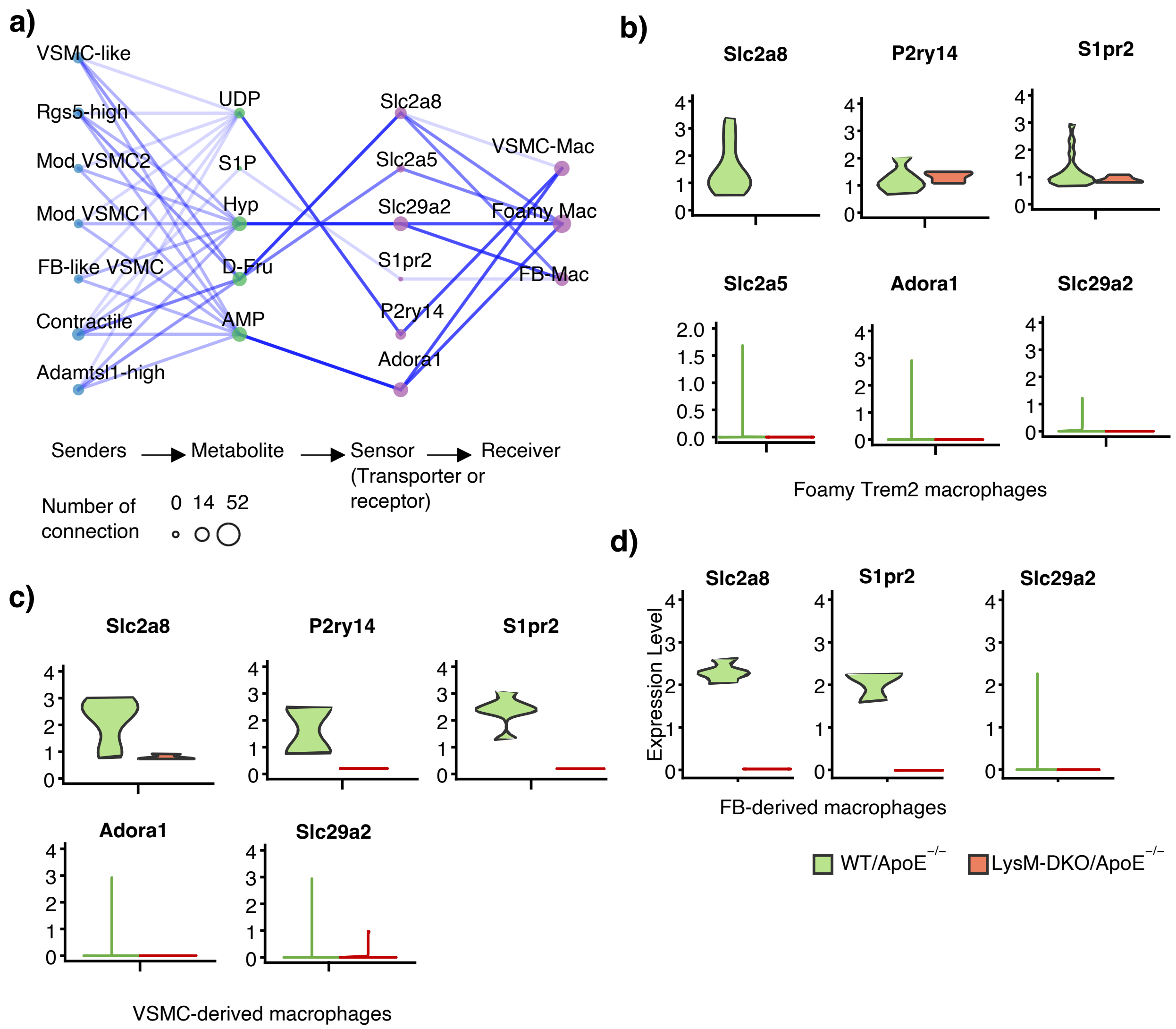
